# Msx genes delineate a novel molecular map of the developing cerebellar neuroepithelium

**DOI:** 10.1101/2023.12.03.569760

**Authors:** Ishita Gupta, Joanna Yeung, Maryam Rahimi-Balaei, Sih-Rong Wu, Daniel Goldowitz

## Abstract

In the early cerebellar primordium, there are two progenitor zones, the ventricular zone (VZ) residing atop the IV^th^ ventricle and the rhombic lip (RL) at the lateral edges of the developing cerebellum. These zones give rise to the several cell types that form the GABAergic and glutamatergic populations of the adult cerebellum, respectively. Recently, an understanding of the molecular compartmentation of these zones has emerged. The *Msx* genes are a family of three transcription factors that are expressed downstream of Bone Morphogenetic Protein (BMP) signaling in these zones. Using fluorescent RNA *in situ* hybridization, we have characterized the *Msx* (Msh Homeobox) genes and demonstrated that their spatiotemporal pattern segregates specific regions in the progenitor regions. *Msx1* and *Msx2* are compartmentalized within the rhombic lip (RL), while *Msx3* is localized within the ventricular zone (VZ). The relationship of the *Msx* genes with an early marker of the glutamatergic lineage, *Atoh1*, was examined in *Atoh1*-null mice and it was found that the expression of *Msx* genes persisted. Importantly, *Msx1* and *Msx3* expressions expanded in response to the elimination of *Atoh1*. These results point to early markers of cerebellar progenitor zones and more importantly to an updated view of the molecular parcellation of the RL with respect to the canonical marker of the RL, *Atoh1*.

## Introduction

During early embryonic development, the neuroepithelium of the cerebellar primordium consists of two primary progenitor zones – the rhombic lip (RL) and the ventricular zone (VZ). The evidence indicates that all glutamatergic cells (glutamatergic cerebellar nuclear neurons, granule cells and unipolar brush cells) arise from the RL while the GABAergic cells (GABAergic cerebellar nuclear neurons, Purkinje cells and interneurons) arise from the VZ (Hoshino et al., 2005; Machold & Fishell, 2005; V. Y. Wang, Rose, & Zoghbi, 2005) (Supplemental Fig. 1). The two progenitor zones are molecularly defined by the non-overlapping expressions of two basic Helix-Loop-Helix (bHLH) transcription factors - *Atoh1* (formerly termed *Math1*) for the RL (Machold & Fishell, 2005; V. Y. Wang et al., 2005) and *Ptf1a* for the VZ (Hoshino et al., 2005). Since these progenitor zones give rise to several cell types over time, it would be important to identify the molecular pathways within the RL and the VZ that play roles in the determination of the different cell types that are generated in the neuroepithelia.

BMP signaling has been studied in cerebellar development and has been shown to be necessary for normal development of both glutamatergic and GABAergic lineages (Alder, Lee, Jessell, & Hatten, 1999; T. C. Ma, Vong, & Kwan, 2020; Qin, Wine-Lee, Ahn, & Crenshaw, 2006; Tong & Kwan, 2013). Activated R-smad is expressed in both the RL and the VZ (Fernandes, Antoine, & Hébert, 2012; Tong & Kwan, 2013). Studies have shown that loss of both BMP signaling components, Smad1 and Smad5 (R-smads), in cerebellum results in defects in RL stem cell specification, loss of nuclear transitory zone (NTZ) and reduced external germinal layer (EGL); and activation of the BMP antagonist NBL1 suppresses RL cell specification (Krizhanovsky & Ben-Arie, 2006; Tong & Kwan, 2013). Overexpressing Smad7 (I-smad that inhibits BMP signaling) in the midbrain-hindbrain boundary (MHB) via *Wnt1*-Cre leads to loss of the choroid plexus and cerebellar morphologic anomalies (Tang, Snider, Firulli, & Conway, 2010). Smad7 (I-smad) is expressed in the EGL suggesting that activated BMP signaling is not required or suppressed in later development processes like EGL formation (Lai, Klisch, Roberts, Zoghbi, & Johnson, 2011). Involvement of BMP signaling in the VZ lineages has had more limited exploration. While the loss of *Smad4* (Co-smad) does not affect the glutamatergic lineage, *En1*-Cre knock-out of *Smad4* results in reduced number of Purkinje cells (Zhou et al., 2003). At an earlier age of E11.5, conditional knock-out of *Smad4* using *En1*-Cre significantly reduces the proliferative KI67-positive VZ progenitors (Fernandes et al., 2012). Recently, a study by Ma et al. (2020) has shown that the gradual spatiotemporal decline in the BMP/Smad gradient across the dorso-ventral axis of the VZ directs the identity transition of the VZ progenitor cells from Olig2-positive Purkinje neuron progenitors to Gsx1-positive interneuron progenitors (T. C. Ma et al., 2020). While it is clear that BMP signaling is important to the developing cerebellum, it is not clear what are the downstream genetic and transcriptional changes that mediate this signaling in the cerebellum, particularly in the progenitor zones of the RL and VZ.

The *Msx* (Msh Homeobox) genes are directly activated by BMP signaling in mice and are suitable candidates for mediating BMP signaling in cerebellar development (Suzuki, Ueno, & Hemmati-Brivanlou, 1997; Takahashi et al., 1998). These homeobox-containing genes are known transcriptional repressors (K M Catron et al., 1995; Katrina M. Catron, Wang, Hu, Shen, & Abate-Shen, 1996; Newberry, Latifi, Battaile, & Towler, 1997; H. Zhang, Catron, & Abate-Shen, 1996). The mouse family consists of three members - Msx1, Msx2 and Msx3. These 3 genes share 98% sequence similarity in their homeodomains (Ekker et al., 1997). Mouse Msx3 and the putative human ortholog VENTX (based on NCBI’s Eukaryotic Genome Annotation pipeline) do not share sequence homology, hinting at a species-based differences in functions and redundancy of the Msx family. In mice, Msx1 and Msx2 have been extensively studied in the context of craniofacial morphogenesis and limb organogenesis (Foerst-Potts & Sadler, 1997; Satokata et al., 2000; Satokata & Maas, 1994; Wu et al., 2003); while in neural development they are known to be expressed in overlapping patterns in many regions including roof plate cells and the adjacent neural tube (Duval et al., 2014; Sunkin et al., 2013; Tríbulo, Aybar, Nguyen, Mullins, & Mayor, 2003). Msx3 has been relatively less studied in the context of development although it is exclusively expressed only in the dorsal CNS in mice, particularly the developing spinal cord and the cerebellum (Sunkin et al., 2013; W. Wang, Chen, Xu, & Lufkin, 1996). Based on the expression of the *Msx* genes in the E9.5-E10.5 murine neural tube, along with strong expression in the neural plate of cephalochordates (Sharman, Shimeld, & Holland, 1999) and ascidians (L. Ma et al., 1996), this family of genes seems to have a strong conserved function in dorsal neural tube patterning.

We had found that all 3 *Msx* genes are expressed early in our cerebellar time-course FANTOM5 transcriptome dataset (Arner et al., 2015; Ha et al., 2019). From the perspective of an importance of BMP signaling in the early developing cerebellum and that the Msx genes could be mediators of this signaling with the early specific expression in the cerebellum, we sought to explore possible roles of this family of genes in the context of mouse cerebellar development.

## Results

### Msx genes are expressed only in the progenitor zones at early time-points

As a first step towards understanding the temporal expression patterns of the *Msx* transcription factors (TFs), we examined the FANTOM5 time-course transcriptome for the developing cerebellum (Ha et al., 2019). All three Msx TFs have a highly dynamic temporal expression signature, with a steep decline after E11.5-E12.5 (Fig 1 a-c). The peak times of *Msx1* and *Msx3* expression are at the early embryonic stages. *Msx2* has a biphasic expression pattern, showing expression at the early embryonic time and an early postnatal expression.

**Figure 1.**
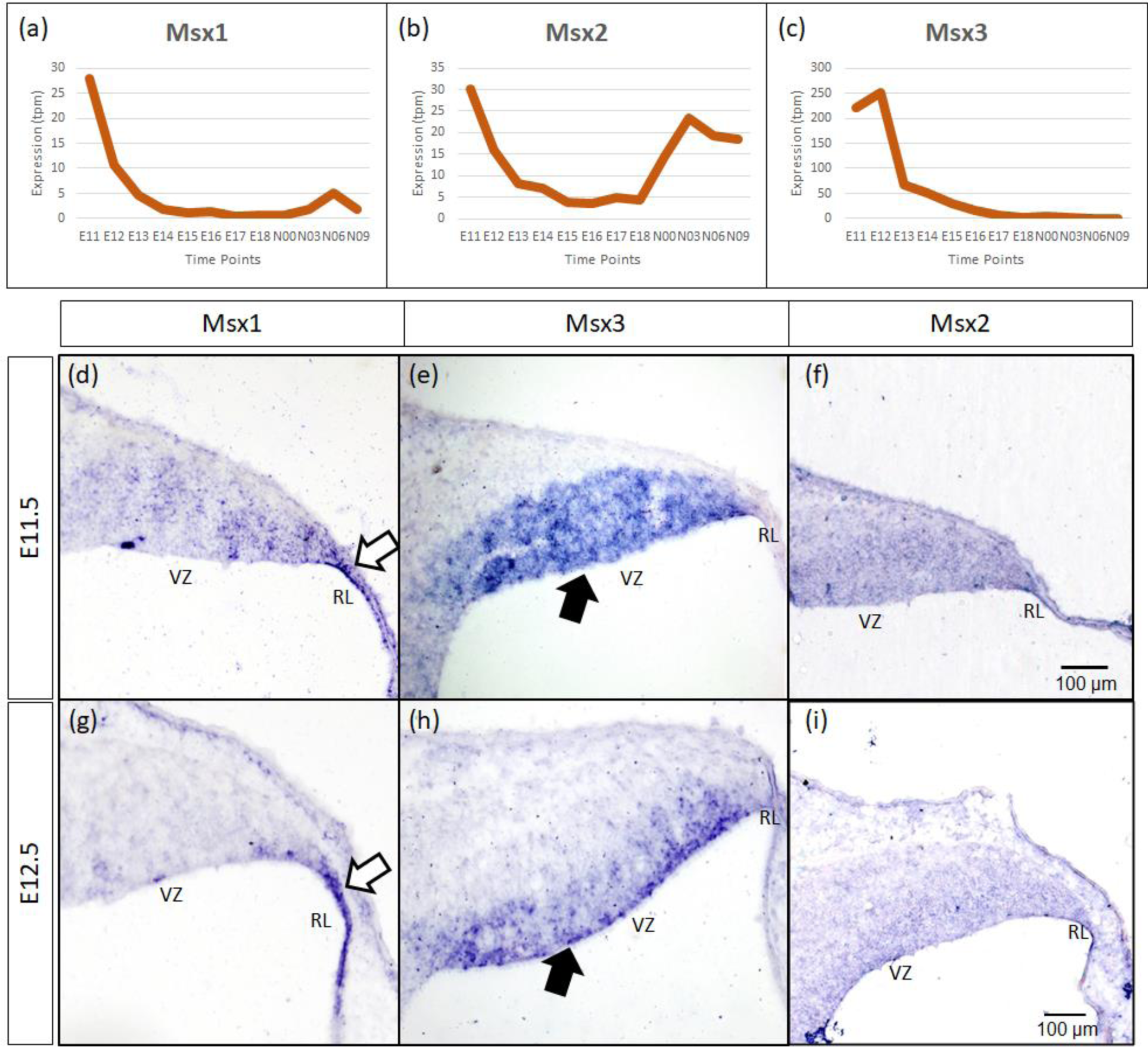
Temporal and spatial expression of Msx genes in the developing cerebellum. (a-c) Graphs show the dynamic nature of Msx expression in the cerebellum across 12 developmental time-points as observed from the RIKEN FANTOM5 transcriptome time-course data. (d-i) Sagittal sections of the RL with the right-side of panels denoting posterior and the bottom-side denoting ventral, with RNA *in situ* hybridization showing Msx genes expressed in the progenitor zones in (d-f) E11.5 and (g-i) E12.5. Msx1 expression is limited to the RL (white arrows in d, g) whereas Msx3 is limited to the VZ (black arrows in e, h). Msx2 expression is detected in the neuroepithelium but the boundary of the expression is not clear (f, i). See supplementary Figure 2 for negative control staining. RL, Rhombic Lip; VZ, Ventricular Zone. Scale bar, 100 µm.

To evaluate spatial expression of the *Msx* genes, we used chromogenic RNA *in situ* hybridization (ISH) to probe for the mRNA on sagittal sections of the developing cerebellum (see Methods for probe and tissue details). *Msx1* is expressed in the rhombic lip (RL) at both E11.5 and E12.5 (white arrows in Fig.1 d, g). In a complementary manner, *Msx3* expression is reserved to the ventricular zone (VZ) (black arrows in Fig.1 e, h). With this *in situ* method, Msx2 expression is found throughout the neuroepithelium, but the boundaries of its expression are less clear (Fig.1 f, i). Expressions of *Msx1* and *Msx3* get more restricted with finer defined boundaries at E12.5 compared to E11.5. Thus, at these early ages, the expressions of all the *Msx* genes are concentrated in the progenitor zones of the neuroepithelium and are absent from the rest of the cerebellar primordium.

### Msx1 and Msx2 expressing cells are compartmentalized within the rhombic lip at E12.5

As indicated in the Introduction, *Msx1* and *Msx2* have been shown to have overlapping expression domains outside of the cerebellum, with sometimes similar and/or redundant functions. Our chromogenic ISH demonstrated that both *Msx1* and *Msx2* are expressed in the cerebellar RL. To further examine their relationship in the cerebellar RL, RNAscope fluorescent *in situ* hybridization (FISH) multiplex assay (F. Wang et al., 2012) was used to double-label *Msx1* and *Msx2* mRNAs; also in regard to *Atoh1*. Interestingly, both *Msx1* and *Msx2* are expressed in cells within the RL but they are expressed in different populations of cells (Figure 2a). At E12.5 *Msx1* is expressed most strongly in the posterior-most tip of the RL that is *Atoh1* negative (white arrows in Fig. 2a, b, d) with much weaker expression in the rest of the RL. This is also observed at E11.5 (Supplementary Figure 4). The same observation of non-overlapping Msx1 and Atoh1 expression is reproduced using antibodies targeting Msx1 and Atoh1 with immunofluorescence (Supplementary Figure 5a-c). *Msx2* expression is within the *Atoh1* expression domain and is not expressed in the *Atoh1*-negative *Msx1*-positive compartment (Figure 2a, c). Thus, *Msx1* and *Msx2* expression regions are non-overlapping and form an *Msx1-Msx2-Msx1* banding pattern (Figure 2c). At postnatal ages, *Msx2*, and not *Msx1*, expression is detected in the granule cells (Supplementary Figure 6c,d). The schematic illustration of this compartmentation at E12.5 is shown in Figure 2g.

**Figure 2.**
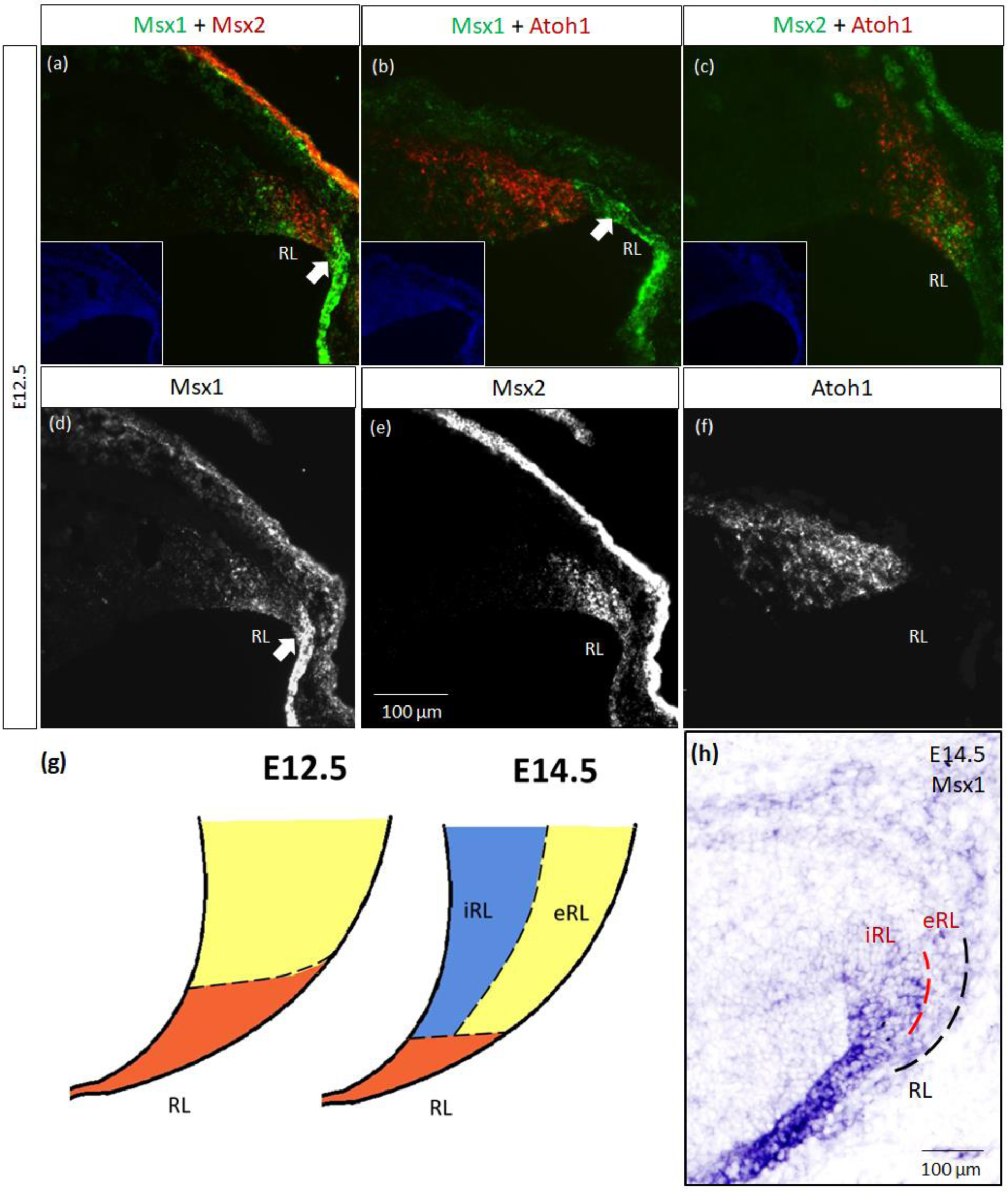
Msx1 and Msx2 expression relative to Atoh1 in the early cerebellum. (a-f, h) Sagittal sections of the RL with the right-side of panels denoting posterior and the bottom-side denoting ventral. (a-c) RNAscope fluorescent RNA *in situ* hybridization (FISH) double-label on sagittal sections of E12.5 cerebellum. (a) Msx1 (green) is expressed highest in the distal tip of the RL (white arrow) posterior to Atoh1 (red). This compartment is Atoh1-negative (b) and maps to the red region shown in (g) at E12.5. (c) Msx2 (green) and Atoh1 (red) are largely overlapping in their expression regions. Msx1 (green) and Msx2 (red) expression regions form an alternative banding pattern and do not overlap with each other. (a-c) Inset shows DAPI (blue) counterstain of the respective cerebellar tissue sections. Roof plate epithelium auto-fluoresces with the fluorescent dyes. (d-f) shows the expression of Msx1, Msx2 and Atoh1, respectively. (g) Schematic illustrating the compartments within the RL at E12.5 and E14.5 based on results by Yeung et al. (2014). At both ages, red represents Wls-positive, Msx1-positive and Atoh1-negative, yellow represents Atoh1-positive, Wls-negative, Msx1-negative. At E14.5 the blue iRL region is Wls-positive and Atoh1-negative. (h) RNA In situ hybridization of Msx1 on sagittal E14.5 section shows stronger expression in the Wls-positive iRL than the eRL. Refer to Appendix Figures 2 and 3 for negative control staining. eRL, exterior RL; iRL, interior RL; RL, Rhombic Lip. Scale bars, 100 µm.

Later at E14.5, chromogenic RNA *in situ* hybridization reveals that *Msx1* expression is most obvious in the iRL (interior RL) compartment (Figure 2h), a region defined by high Wls expression and low Atoh1 expression (Yeung et al. 2014). Immunofluorescence (IF) double labeling with Msx1 and Atoh1 antibodies detected Msx1 expression in the iRL but no expression of Atoh1 (Supplementary Figure 5d-f). At this age Atoh1 expression is restricted to the eRL (exterior RL); the two proteins do not overlap in the RL at E14.5. The schematic of this compartmentation at E14.5 is illustrated in Figure 2g.

The observation that Msx1 expression is highly restricted in the E14.5 iRL prompted us to examine its relationship with Wls, a known marker of the iRL (Yeung et al., 2014). To this end, we examined Msx1 expression in the *Wls*-reporter mouse strain that expresses ꞵgal under the *Wls* promoter at both E12.5 and E14.5 (Yeung et al., 2014). Double labeling of Msx1 and ꞵgal in E12.5 *Wls*-reporter cerebellum demonstrated an overlapping expression of both molecules in the tip of the RL and the rest of the RL area (Red arrows in Figure 3). Msx1 expression, however, is absent in the stream of ꞵgal-expressing cells that emanate from the RL and travel into the subpial stream (White arrows in Figure 3). In the E14.5 *Wls*-reporter cerebellum, Msx1 and ꞵgal continue to co-express in the iRL (Yellow arrows in Figure 3), while ꞵgal still marks the cells that migrate out of the RL into the EGL; Msx1 is absent from the eRL and EGL (Figure 3).

**Figure 3.**
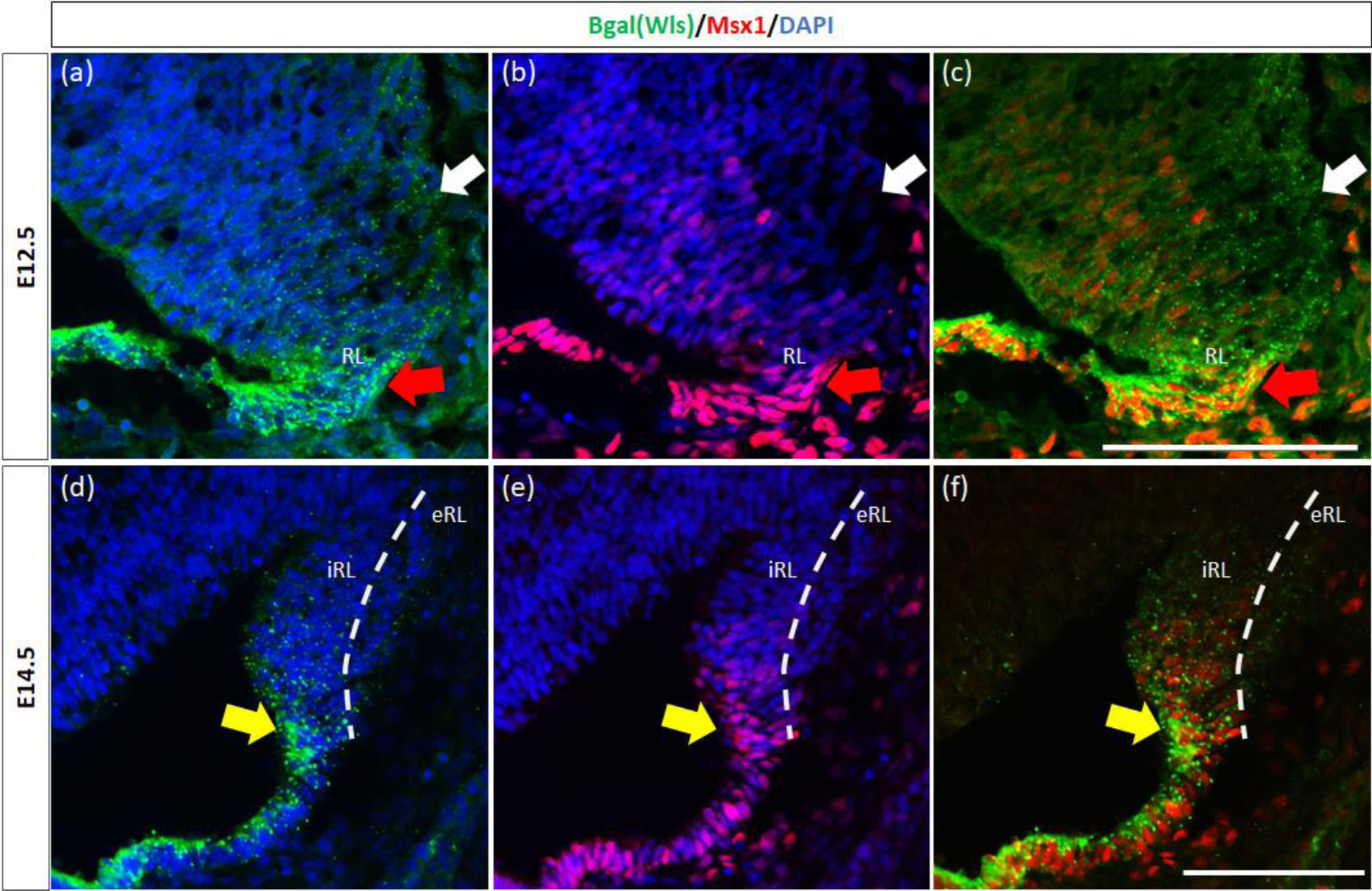
Msx1 and Wls expression in the developing cerebellar rhombic lip. Expression of Wls and Msx1 was examined in Wls-βgal reporter mouse embryos at E12.5 (a-c) and E14.5 (d-f). At E12.5, Msx1 and Wls are co-expressed strongly in the distal tip of the RL (red arrows), co-expression of both molecules can also be seen in the RL area. Msx1, however, is not expressed in the Wls-positive subpial stream (white arrows) populated by the RL-lineage progenitors emanating from the RL. At E14.5, Msx1 and Wls continue to co-express and localize to the iRL (yellow arrows) while Msx1 expression is absent in the eRL. eRL, exterior face of the RL; RL, Rhombic Lip; iRL, interior face of the RL. Scale bars, 100 µm.

Previously, Duval et al. (2014) have shown through lineage tracing analysis of Msx1 and Msx2 in the murine dorsal spinal cord that almost all Atoh1-positive cells at E10.5 arise from progenitors expressing Msx1 as early as E9.25 (Duval et al., 2014). As Msx1 is expressed in the same compartments as Wls in the RL, this expression pattern of Msx1 also supports the possible upstream regulatory role of Msx1 in relation to Atoh1 in the cerebellum. To investigate the relationship of Msx1, as well as Msx2, with Atoh1, we examined their expression in the *Atoh1*-null cerebellum. In the E12.5 *Atoh1*-null cerebellum, *Msx1* expression in the RL persisted (Figure 4). Moreover, *Msx1* expression is no longer restricted to the distal tip of the RL and expands to a larger domain in *Atoh1*-null RL (Figure 4). *Msx2* expression is also found to persist in the *Atoh1*-null cerebellum, and its expression pattern is similar to that in the control cerebellum (Figure 4). While expression of *Msx1* and *Msx2* persisted in *Atoh1*-null cerebellum, the *Msx1-Msx2-Msx1* banding pattern we previously observed in the wildtype RL is absent in the *Atoh1*-null.

**Figure 4:**
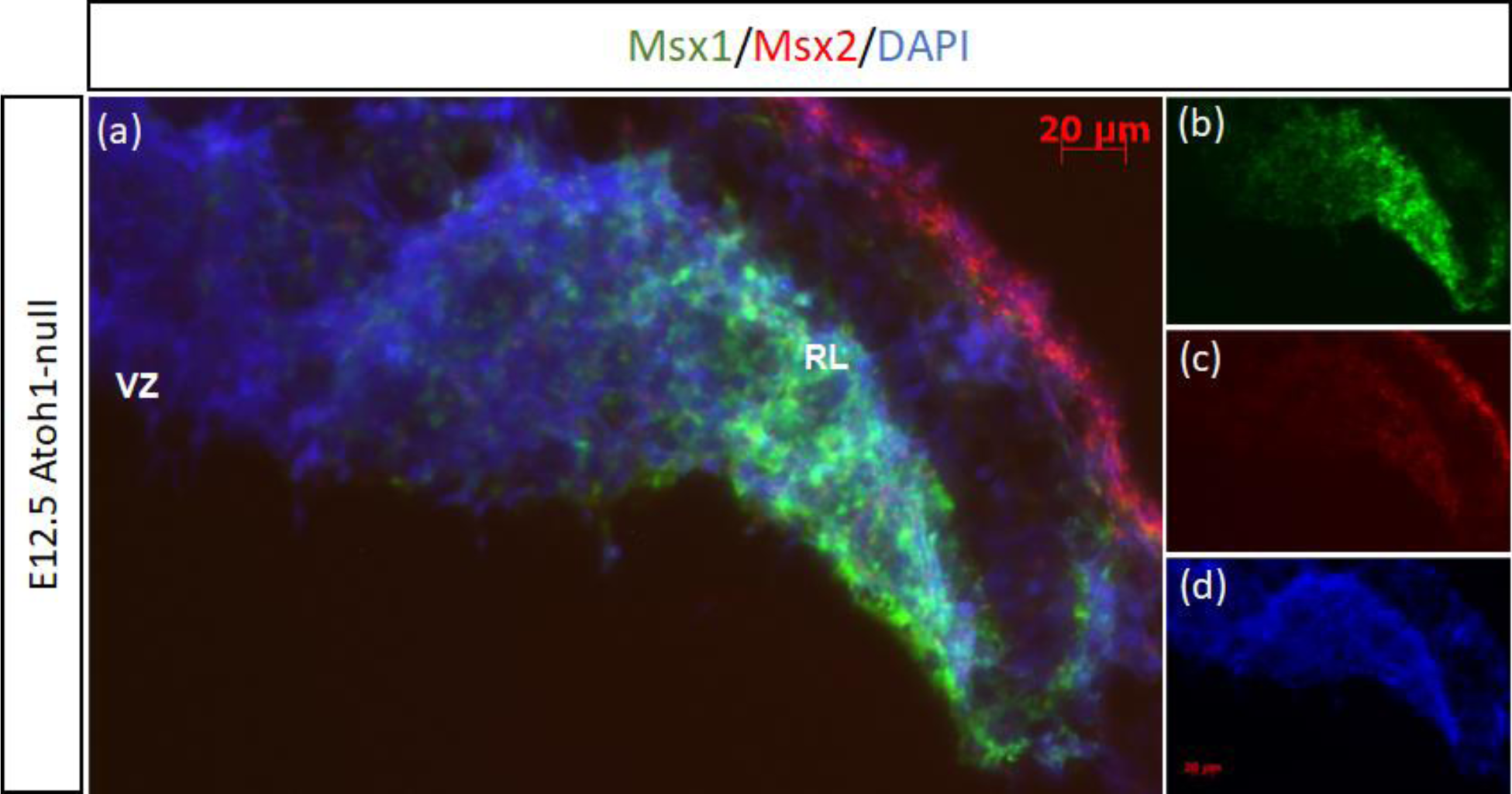
Msx1 and Msx2 expression in E12.5 Atoh1-null cerebellum. (a-d) The FISH double-labeling of Msx1 and Msx2 in the E12.5 *Atoh1*-null cerebellum indicates their expression persistence (a). While the expression pattern of Msx1 expands to a larger domain in *Atoh1*-null RL (b), Msx2 expression is similar to that in the control cerebellum (c). (d) is DAPI (blue) as counterstain. RL, Rhombic Lip; VZ, ventricular zone. Scale bars, 20 µm.

Similar to our current results of Msx1 and Msx2 being upstream of Atoh1 in the cerebellum, it is known from our previous study that Wls in the cerebellum is upstream of Atoh1 (Yeung et al., 2014). To determine the relationship between Msx1 and Wls, we examined *Msx1* expression in the *Wls* knockout cerebellum. Figure 5 illustrates that in the E12.5 *Wls* knockout cerebellum, *Msx1* expression is still observed in the RL, suggesting that Msx1 expression in the cerebellum is independent of Wls.

**Figure 5.**
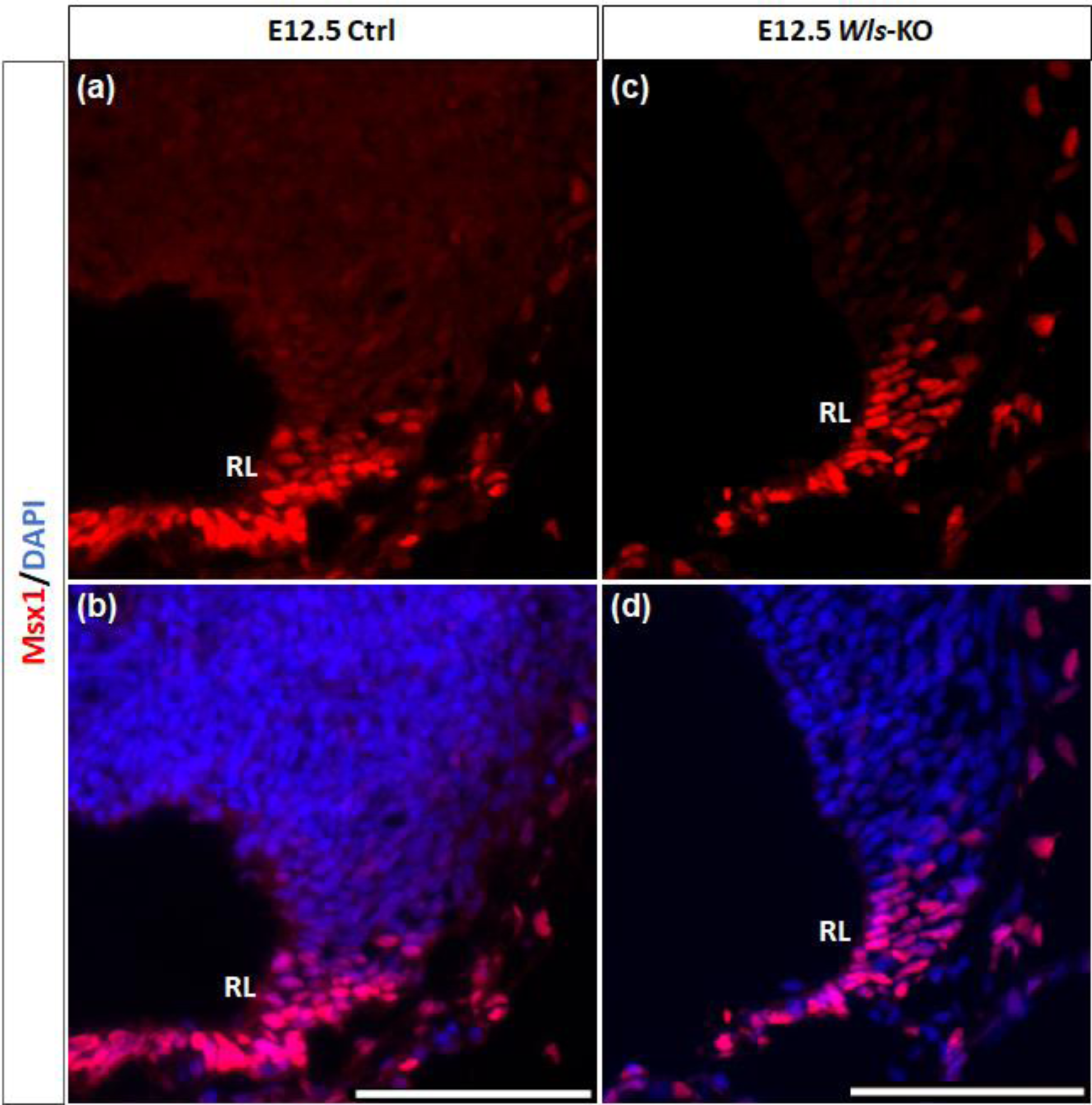
Msx1 expression in E12.5 *Wls*-knockout mutant and control cerebella. In control cerebellum (a, b), Msx1 is expressed in the RL. Expression of Msx1 in the RL is persisted in the *Wls* knockout cerebellum (c, d) and similar to Msx1 expression in the control RL. RL, rhombic lip. Scale bars, 100 µm.

### Msx3-positive cells mark the boundary region in the neuroepithelium between the Atoh1 and Ptf1a domains at E12.5

As seen with chromogenic ISH analysis, *Msx3* is expressed throughout the VZ at E11.5 and E12.5 (Black arrows in Figure 1e, h). To determine if *Msx3* expression extends to the RL, double-labeling FISH for *Msx3* and the RL marker *Atoh1* was examined at E12.5 (Figure 6). This revealed that *Msx3* expressing cells do not overlap with *Atoh1* expressing cells and a boundary can be observed between their respective domains (Figure 6a-c). Double-labeling FISH of *Msx3* with *Ptf1a* at the same age revealed that they largely overlap in their expression domains in the VZ, with a notable exception that their posterior-most expression boundaries, near the RL, do not coincide (Figure 6d-f). This is also observed at E11.5 (Figure 6g-i). *Msx3* expression extends more posteriorly compared to *Ptf1a* expression at E11.5 and E12.5 (the region between the white arrows in Figure 6d, g). *Msx3* expression at its posterior edge creates a molecular demarcation between the non-overlapping *Atoh1* and *Ptf1a* expressing regions. To test if the posterior-edge expression of *Msx3* is restricted by the expression of *Atoh1*, we examined *Msx3* expression in the absence of *Atoh1*. In E12.5 *Atoh1*-null cerebellum, cells expressing Msx3 were no longer restricted to the VZ with an expansion of expression into the rhombic lip (Figure 7). Despite the overlapping expansion of *Msx1*+ and *Msx3*+ cells, no cells are found to co-express both genes in the E12.5 *Atoh1*-null cerebellum (Figure 7a and c-f).

**Figure 6.**
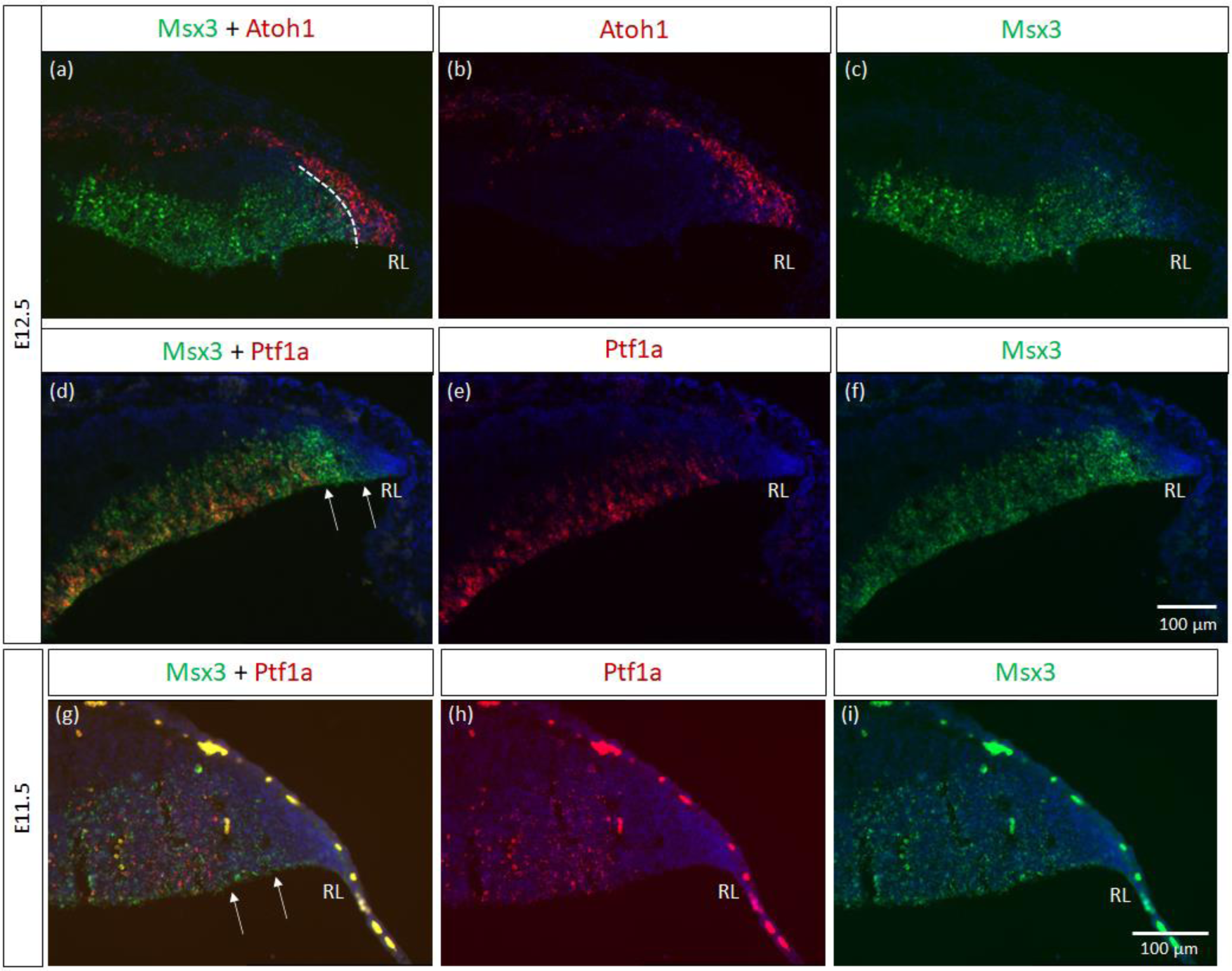
Msx3 expression relative to Atoh1 and Ptf1a in the early cerebellum. (a-f) RNAscope FISH double-label on E12.5 cerebellum. (a-c) Co-labeling of Msx3 (green) and Atoh1 (red) illustrates that Msx3 and Atoh1 do not overlap; this boundary is marked by the dashed line. (d-f) Co-labelling of Msx3 (green) and Ptf1a (red) illustrates a large overlap in their regions of expression in the VZ. The Msx3 boundary extends further than the Ptf1a boundary and abuts the RL (arrows). (g-i) RNAscope double-label on sagittal E11.5 sections with Msx3 (green) and Ptf1a (red) shows the same Msx3-exclusive region near the RL (arrows). Note that the roof plate epithelium has auto-fluorescence giving rise to the blob-like artefacts (refer to Supplementary Figure 5a for negative control staining). All panels have DAPI (blue) as counterstain. RL, Rhombic Lip. Scale bars, 100 µm.

**Figure 7.**
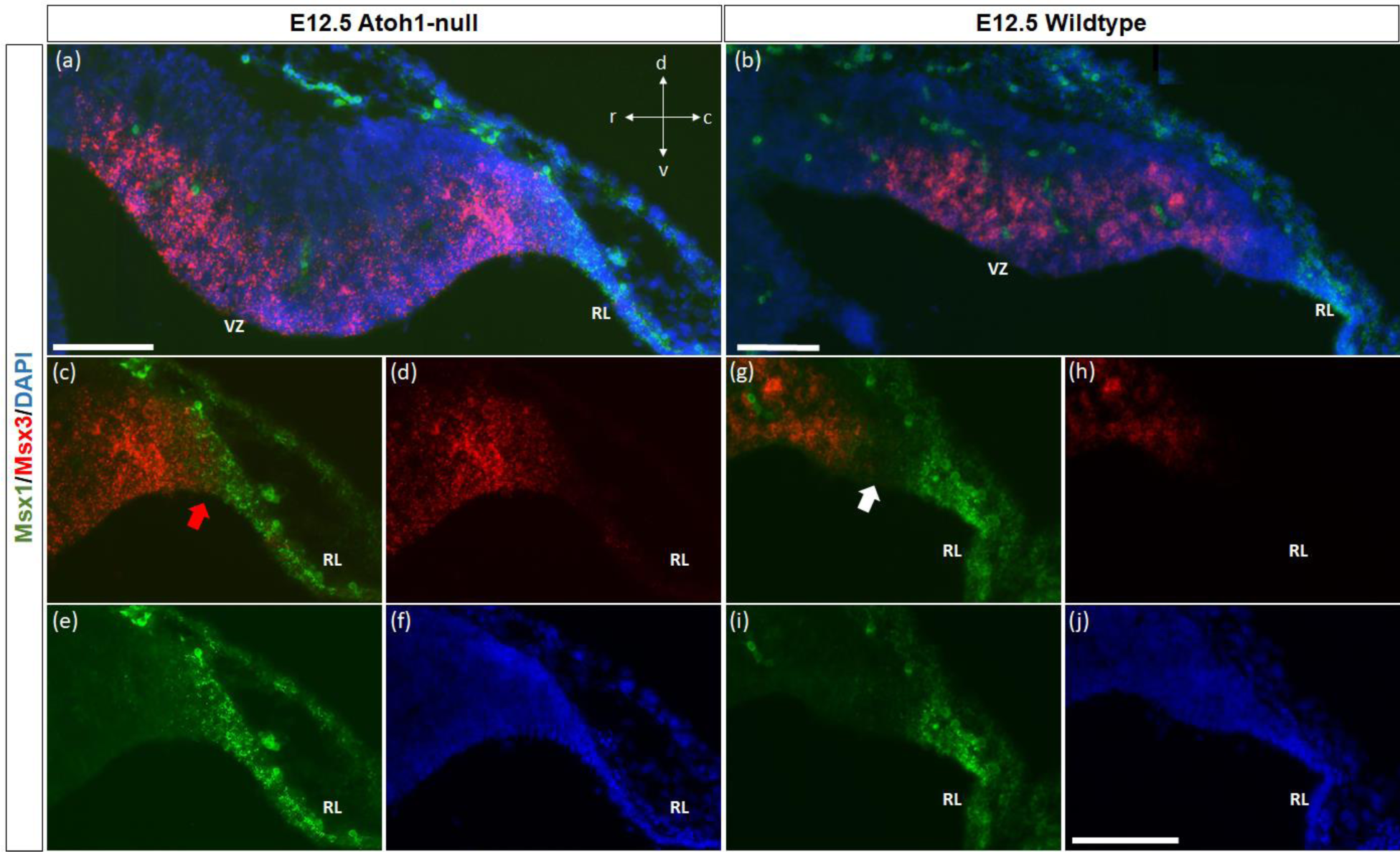
Msx1 and Msx3 expression in E12.5 Atoh1-null and wildtype cerebella. (a-b) The FISH double-labeling of Msx1 and Msx3 in the E12.5 *Atoh1*-null (a) and wildtype cerebella (b). (c-f) Expression of Msx1 and Msx3 expands and overlaps in *Atoh1*-null RL as denoted by the red arrow in (c). Panels (a, c-f) indicate Msx1 and Msx3 expression persistence in the *Atoh1*-null cerebellum. (a), this does not result in cells that co-express Msx1 and Msx3 (a, c-e). (g-i) A very different picture is shown in the wildtype cerebellum where there is a notable gap between cells that express Msx3 and Msx1 as noted by white arrow in (g). (f,j) DAPI (blue) used as a counterstain. RL, Rhombic Lip; VZ, ventricular zone. All scale bars, 100 µm.

### Msx3-positive cells are restricted to the posterior region of the proliferative ventricular zone at E14.5 in the lateral cerebellum

While *Msx3* is expressed throughout the VZ at E11.5 and E12.5, *Msx3* expression becomes restricted within the VZ at later time-points. Interestingly the *Msx3* expression is spatially dynamic along the lateral-medial axis at E14.5 (Figure 8). In the most lateral aspect of the cerebellum, *Msx3* occupies a small region in the posterior part of the VZ near the RL (Figure 8a,b) and progressively occupies the entire VZ at the most medial aspect of the cerebellum (Figure 8e,f).

**Figure 8.**
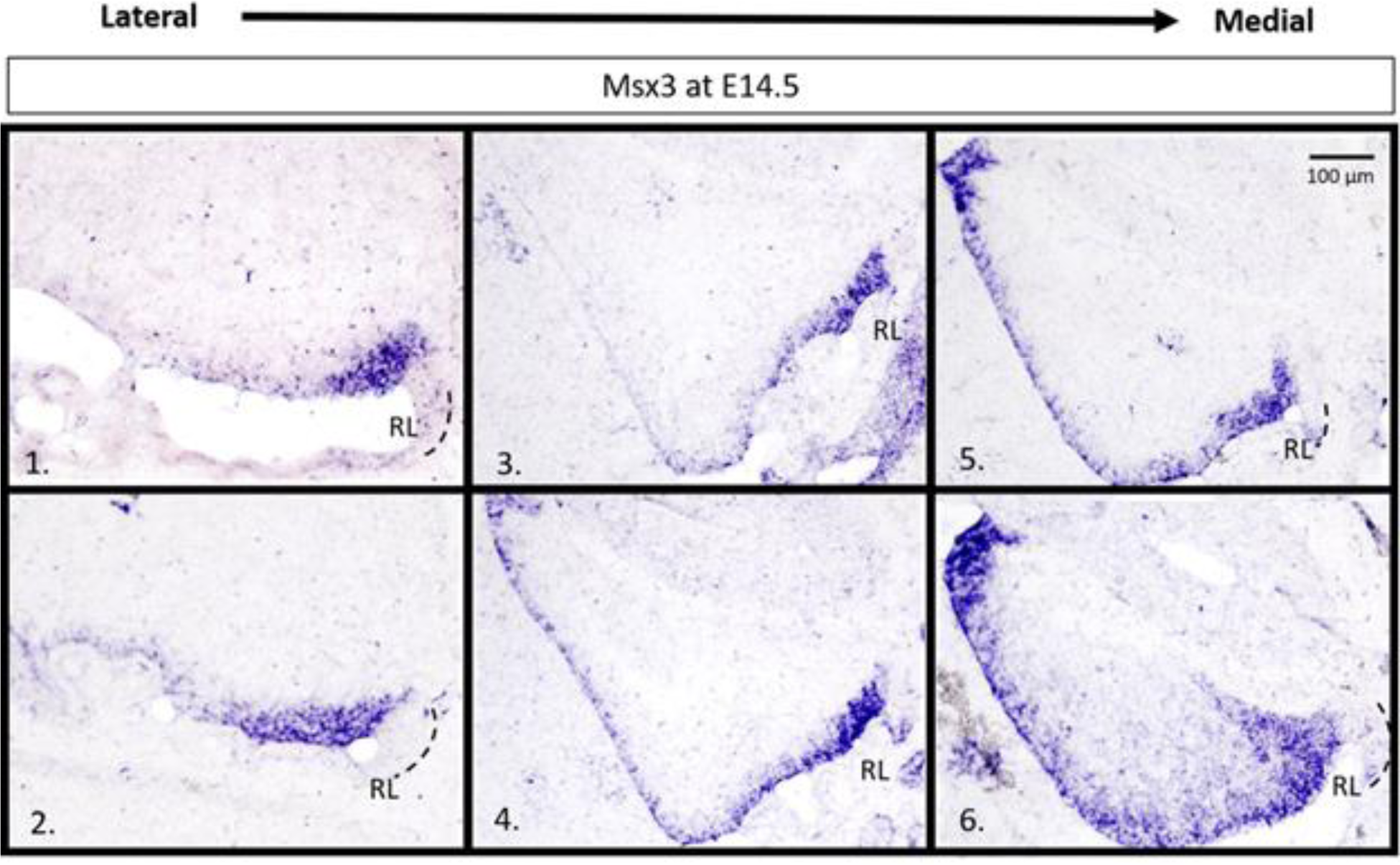
*Msx3* expression is spatially dynamic within the VZ at E14.5. (a-f) are sagittal sections of wildtype cerebellum at E14.5, with the right side of the panels denoting posterior and bottom side denoting ventral. RNA in situ hybridization of Msx3 in increasing order of relative lateral-medial positions with (a) being the most lateral, (b) being more medial than (a), and so on with (f) being the most medial. Msx3 gets restricted to the posterior end of the lateral VZ (a,b) and progressively occupies the entire VZ in the medial sections (e,f). Refer to Appendix Figure 2c for negative control staining. RL, Rhombic Lip. Scale bar, 100 µm.

We examined cell proliferation at the VZ within the *Msx3* expressing domain using FISH along with BrdU immunohistochemistry. We used BrdU labeling to mark cells that are proliferating one hour before we harvest the embryos at E14.5 (Figure 9). *Msx3* expression along the lateral-medial axis was found to be strictly within the proliferative region of the neuroepithelium.

**Figure 9.**
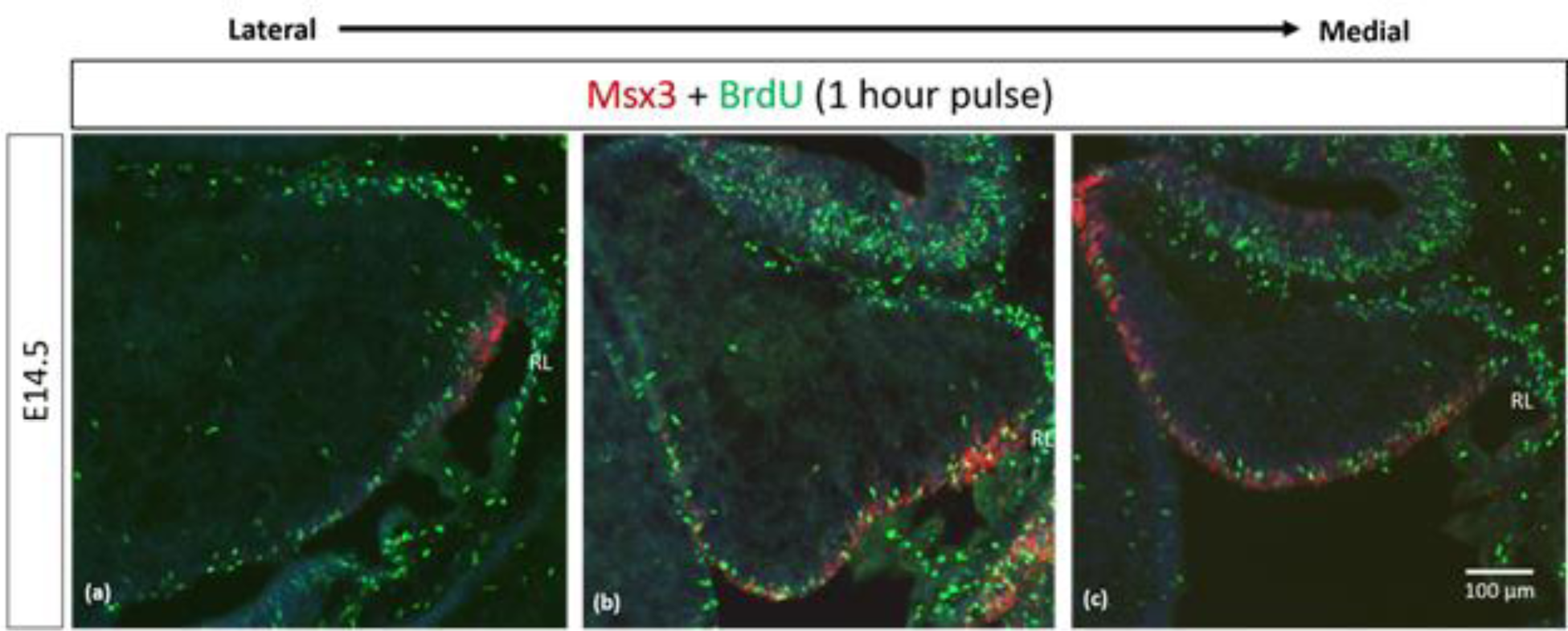
Msx3 is expressed strictly within the proliferative neuroepithelium. (a-c) are sagittal sections of wildtype cerebellum at E14.5, with the right side of the panels denoting posterior and bottom side denoting ventral. RNAscope FISH of Msx3 (red) with fluorescence immunohistochemistry of BrdU (green) for mouse pulsed with BrdU one hour before E14.5 embryos were harvested. Relatively (a) is the most lateral and (c) is the most medial. RL, Rhombic Lip. Scale bar, 100 µm.

## Discussion

The members of the *Msx* gene family are homeobox-containing, highly conserved transcription factors, previously unexplored in the context of the cerebellum. *Msx* genes are known best as downstream effector molecules of BMP signaling, which is a crucial signaling pathway for the cerebellum via the roofplate epithelial cells posteriorly adjacent to the developing cerebellar primordium (Krizhanovsky & Ben-Arie, 2006). In our examination, the *Msx* genes are found to pattern the proliferative neuroepithelium of the early embryonic cerebellar primordium. High resolution RNAscope reveals that *Msx1* marks the *Atoh1*-negative distal tip of the RL, while *Msx2* does not overlap with *Msx1* and marks the *Atoh1*-positive rhombic lip; *Msx3* is expressed in the *Ptf1a*-positive ventricular zone but also demarcates a *Ptf1a*-negative region that neighbors the rhombic lip. This spatial patterning brings new dimensions to our understanding of compartmentation of the cerebellar neuroepithelium and is summarized in Figure 10a.

**Figure 10.**
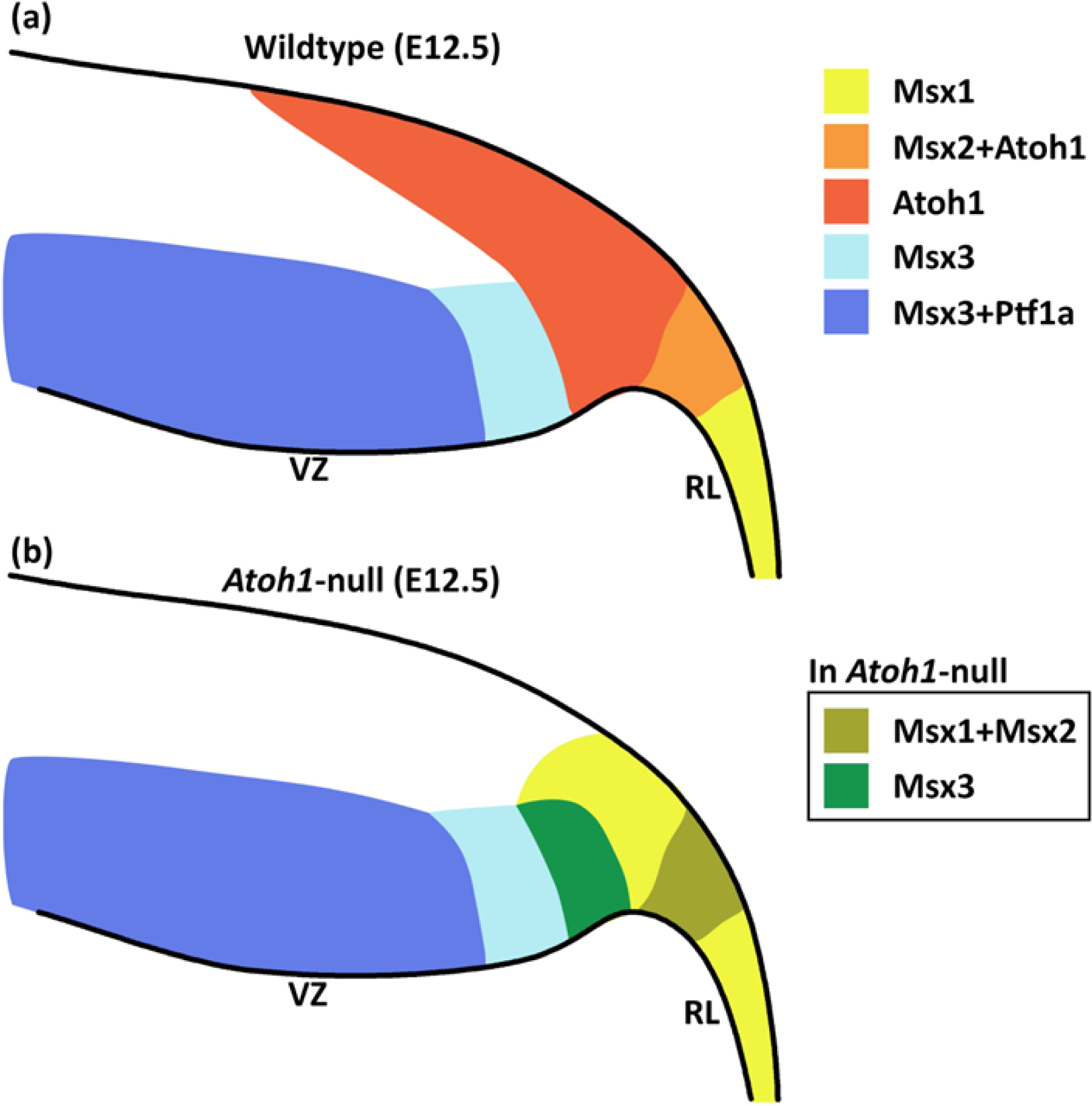
Schematic illustration of Msx expression in the early embryonic cerebellar neuroepithelium. (a) During early cerebellum development, expression patterns of Msx genes within the neuroepithelium demarcate the VZ and RL into molecularly distinct regions. Msx1 expression marks the distal tip of the RL that is Atoh1-negative. Msx2 is found to be co-expressed with Atoh1 in the RL. Msx3 is expressed in the VZ and largely overlapped with expression of Ptf1a, although Msx3 expression extends beyond the posterior end of the Ptf1-positive zone and abuts the Atoh1-positive RL. (b) In the Atoh1-null cerebellum, all three Msx genes expression persisted. The expression of Msx1 and Msx3 in the absence of Atoh1 is observed to expand into the presumptive Atoh1 region in the RL. RL, rhombic lip; VZ, ventricular zone.

We find that all 3 *Msx* genes persist in the *Atoh1*-null cerebellum, placing them as BMP signaling effector molecules upstream (*Msx1* and *Msx2*), or independent (*Msx3*), of *Atoh1*. Upon *Atoh1* ablation, *Msx1* expression extends into the region formerly colonized by *Atoh1*-positive cells of the rhombic lip. *Msx3* expression extends posteriorly towards the rhombic lip. These findings show that *Atoh1* inhibits the expression of *Msx1* and *Msx3*, placing the *Msx* genes upstream to, and in a dynamic regulatory network with, *Atoh1* (see Figure 10b for Summary Schematic). As external signaling molecules, the BMPs have been implicated in the specification of cerebellar cell types but their downstream molecular cascades are unknown. The results of this study present the *Msx* genes as strong candidates facilitating this BMP signaling in cerebellum development.

### Msx3 as a key regulator of the ventricular zone

*Msx3* is the most understudied member of the family. *Msx3* is yet to be confirmed as a direct transcriptional target of BMP signaling, although ectopic BMP expression can induce *Msx3* (Shimeld, McKay, & Sharpe, 1996). Unlike the other family members, *Msx3* is present exclusively in the dorsal murine CNS tissue (Sunkin et al., 2013). Our characterization of *Msx3* at E12.5 with known germinal zone markers, Ptf1a and Atoh1, revealed an intriguing pattern along the VZ and RL at this time of early cerebellar development. The current view of the cerebellar neuroepithelia depicts the VZ as marked by Ptf1a+ cells throughout. In our present findings, however, *Msx3* is expressed in the VZ and forms a boundary with the *Atoh1*+ cells in the RL. While *Msx3* largely overlaps with *Ptf1a* in the VZ, Msx3 expression extends beyond the posterior end of Ptf1a-positive zone, as shown by a gap between *Ptf1a*-positive VZ and *Atoh1*-positive RL; and here there is an *Msx3*-exclusive region. The *Msx3*-exclusive region, defined as *Atoh1*-negative/*Ptf1a*-negative/*Msx3*-positive, has not been previously identified and invites the question of what cells arise from it.

In a recent single-cell RNA sequencing of the early cerebellum, a rare group of cells with mixed features of RL and VZ (i.e. expressing both *Atoh1* and *Ascl1*) have been identified suggesting that parts of the RL and VZ lineages may have a common origin (Khouri-Farah, Guo, Morgan, Shin, & Li, 2022). Zhang et al (2021) report a common progenitor that generates glutamatergic and GABAergic lineages, and also detected a similar rare cell group (Atoh1- and Ptf1a-positive) at the border of RL and VZ (T. Zhang et al., 2021). This seems to overlap with the *Msx3*-exclusive region we identify in the present study. Further research that examines the lineage tracing of *Msx3*-expressing cells (possible bipotent progenitors) in comparison to that of *Ptf1a*-expressing cells (GABAergic lineage) will address whether *Msx3* demarcates a population of bipotential progenitors in the VZ of cerebellar primordia.

*Msx3* expression gets more restrictive at E14.5, receding to the posterior most part of the ventricular zone, which would be exclusively the *Atoh1*-negative/*Ptf1a*-negative/*Msx3*-positive region as described before. This receding expression pattern in the ventricular zone is also shown by Olig2, a transcription factor that specifies the Purkinje cell progenitors, as well as the canonical BMP and p-SMAD1/5 molecules that form a morphogenetic gradient (T. C. Ma et al., 2020; Seto et al., 2014). Another transcription factor crucial to this zone is Gsx1 that specifies the interneuron progenitors (Seto et al., 2014). As Purkinje cells are born first from the ventricular zone, followed by interneurons, the progenitor cells of this zone express first Olig2 and then Gsx1 (Seto et al., 2014). Recently, Ma et al. (2020) have shown that the BMP and p-SMAD1/5 gradient directs the Olig2-Gsx1 based progenitor fate transition in the cerebellar ventricular zone by suppressing Gsx1 expression in the Olig2 domain of the posterior VZ (T. C. Ma et al., 2020). If Msx3 works downstream of BMP signaling to maintain the Olig2 domain, an interesting question is whether Msx3 can suppress interneuron fate or enable Purkinje cell fate, or both? In the study by Liu et al. (2004), a decrease in Pax2 positive interneurons was observed upon Msx3 overexpression in the chick dorsal neural tube (Y. Liu, Helms, & Johnson, 2004). Additionally, in the postnatal mouse cerebellum, *Msx3* expression can be detected in the Purkinje cells but not in the interneurons (Sup. Fig. 7). A key question is whether Msx3 plays a similar role in specifying the cell populations coming from the cerebellar ventricular zone in an anterior-posterior specific pattern.

### Msx1 and 2 as key regulators of the rhombic lip

*Msx1* and *Msx2* are direct transcriptional targets of BMP signaling. Research on the roles of Msx1 and Msx2 in limb and tooth organogenesis points to an association between Msx genes and the extracellular signaling control of the balance between proliferation and differentiation (Dodig et al., 1999; Lallemand et al., 2005; Y.-H. Liu et al., 1999; Nechiporuk & Keating, 2002; Odelberg, Kollhoff, & Keating, 2000; Satokata et al., 2000). Many studies suggest a possible redundancy in Msx1 and Msx2 functions in various tissue types or organ systems (Bei & Maas, 1998; Katrina M. Catron et al., 1996; Chen, Chen, Ishii, Sucov, & Maxson, 2008; J. Han et al., 2007; M. Han, Yang, Farrington, & Muneoka, 2003; Lallemand et al., 2005). In the context of the cerebellum, we noted that in adult stages, postmitotic granule cells express *Msx2* but not *Msx1* (Sup Fig 6). Our high-resolution expression analysis across early embryonic time and space reveals that *Msx1* and *Msx2* are expressed in the rhombic lip, albeit in different regions. *Msx2* is largely overlapping with *Atoh1*, while *Msx1* has the strongest expression in an *Atoh1* negative region of the rhombic lip that is adjacent to the roofplate. Upon *Atoh1* ablation, we found that *Msx2* expression remains unchanged but *Msx1* expressing cells expand. Based on distinct expression patterns of *Msx1* and *Msx2* in early and late stages, we expect their functions to also be distinct in the cerebellum.

In previous work on the developing spinal cord, *Msx1* was also shown to be upstream of *Atoh1* in the dorsal neural tube. Duval et al.’s study of *Msx1-Msx2* double mutant mouse embryos found that the *Atoh1*-expressing dorsal neural tube progenitor cells no longer expressed Atoh1 (Duval et al., 2014). Additionally, they have shown through lineage tracing analysis of Msx1 in the murine dorsal spinal cord that almost all Atoh1-positive cells at E10.5 arise from progenitors expressing Msx1 as early as E9.25. Their conclusion from these findings has been that Msx1 is an activator of Atoh1. It is likely that a similar relationship exists in the cerebellum, and thus we propose Msx1 to be an activator of Atoh1 in the cerebellar rhombic lip as well.

### Msx1 as an early, Atoh1 independent marker of the rhombic lip

Seminal scRNAseq studies on the developing cerebellum identified the presence of *Msx1* in ‘roofplate-like cells’ pointing to a region in the rhombic lip adjacent to the roofplate (Carter et al., 2018; Wizeman, Guo, Wilion, & Li, 2019). We have previously shown that Wls, the main regulator of Wnt molecule secretion, is expressed in cells very similar to the cells that are found to express Msx1 (Yeung & Goldowitz, 2017; Yeung et al., 2014). We also demonstrated an upstream role of Wls relative to Atoh1 by examining the *Atoh1*-null cerebellum. Given that Msx1 and Wls form a distinct boundary with Atoh1, being expressed in the distal-most tip of the RL, we expected Msx1 to be upstream of Atoh1. We confirmed in the *Atoh1*-null cerebellum, that Msx1 expression persisted and expanded. We noted that the former Atoh1-Msx1 boundary in the wildtype RL was blurred due to a spread of Msx1+ cells. Our findings confirm that Msx1 is indeed upstream of Atoh1, and further suggest that Atoh1 has an inhibitory effect on Msx1. Additionally, we found that Msx1 expression persists in a *Wls*-cKO cerebellar RL, showing that Msx1 is independent of WNT signaling.

### Msx genes and the fate of cells in the early cerebellum

Evolution of our understanding of the progenitor cells of the cerebellum has been quite remarkable over the last 50 or so years and highlight the technical advances that have been brought to bear on the analysis of cell lineage in the most complex biological system, the brain. The use of genetically inducible fate mapping has yielded a clear picture of the neurotransmitter-based lineages of the cerebellum: where GABAergic neurons arise from the ventricular neuroepithelium and are marked by Ptf1a (Glasgow, Henke, Macdonald, Wright, & Johnson, 2005; Hoshino et al., 2005) and where glutamatergic neurons arise from the rhombic lip and are marked by Atoh1 (Machold & Fishell, 2005; V. Y. Wang et al., 2005).

The work in our lab has focused on the glutamatergic population that emerges from the rhombic lip. In these studies, we found that the rhombic lip is organized in an inner and outer manner where the inner region is hypothesized to be a more purely generative zone while the outer region is composed of cells whose fate is more restricted. The inner zone (iRL) is marked by cells that express Wls and do not express Atoh1 (Yeung et al., 2014). The outer zone is marked by cells that express Atoh1 with diminished expression of Wls. The current study confirms and provides additional detail to this picture, with Msx1 mapping to the inner region of the RL. The role of Msx1 as a transcriptional repressor of pro-neural and pro-differentiation markers such as Atoh1, Ascl1, Ngn1, Ngn2 and Pax7, as found in the dorsal neural tube (Y. Liu et al., 2004), would fit this model; Msx1 repressing pro-differentiation markers in the cerebellar iRL to keep the progenitor pool in a less-specified state, in tandem with Wls.

Furthermore, with the deletion of *Atoh1*, both *Msx1* and *Msx2* continued to be expressed and in a dynamic manner that indicates that they are upstream to Atoh1 and suggest a regulatory role in cell lineage determination. It is interesting to note that a similar organization of inner and outer regions of the RL has been found in the human RL (Haldipur et al., 2019) and the molecular heterogeneity of these regions may be the key to understanding and treating the most common form on pediatric brain tumor, the medulloblastoma (Hendrikse et al., 2022; Vladoiu et al., 2019).

The present study is the first to examine the patterning and potential roles of the *Msx* genes in cerebellar development. Our results place the *Msx* genes as strong candidates for facilitating BMP signaling in the cerebellum. As transcription factors that are immediate effectors of external BMP signaling from the roof plate, the *Msx* genes are likely to be upstream players of molecular cascades underlying transcriptional regulation of cell types emerging during cerebellar development, given that they play similar roles in limb, tooth, and spinal cord development (Bei & Maas, 1998; Duval et al., 2014; J. Han et al., 2007; Lallemand et al., 2005; Y. Liu et al., 2004). Further studies cementing the regulation of the *Msx* genes by BMP/SMAD signaling in cerebellum would provide directions in this emerging picture of the earliest cell fate decisions in cerebellar development and could likely provide insights into the origin and management of medulloblastoma (Hendrikse et al., 2022; Smith et al., 2022).

Based on our findings, we present the *Msx* genes as candidate novel regulators of early cerebellar progenitors that precede lineage-committed progenitors like the Atoh1-positive cells in the rhombic lip. The early expression patterns of the *Msx* genes suggest a potential function in progenitor cell maintenance and specification, and present as strong candidates for facilitating BMP signaling in cerebellum development.

## Acknowledgements

The authors are grateful to RIKEN FANTOM 5 Consortium for the mammoth technical and informatic efforts applied to the successful time course project and to Huda Zoghbi for the providing of *Atoh1* knockout and control embryos for analysis. The support for trainees was provided by UBC Academic Award (MSc) – Cordula and Gunter Paetzold Fellowship (Ishita Gupta), Michael Smith Health Research BC Trainee Award and BC Children’s Hospital Research Institute Mining for Miracles Postdoctoral Fellowship (Maryam Rahimi-Balaei). This work was supported by the Natural Sciences and Engineering Research Council (NSERC) of Canada, the BCCHRI Brain, Behaviour and Development Theme, and the Howard Hughes Medical Institute (HHMI).

## Methods

### Animal husbandry

The *Wls*-reporter (*Wls^LacZ/+^*) mouse strain was bred and genotyped according to the protocol previously described (Yeung et al., 2014). The *Wls* knockout embryos were generated, phenotype and genotyped according to the protocol previously described (Yeung & Goldowitz, 2017). To harvest tissues at embryonic time points, timed pregnancies were set up. The morning a vaginal plug was detected was designated as embryonic day 0.5 (E0.5). All animal procedures are conducted in accordance with the guidelines of the Animal Care Committee (ACC) at University of British Columbia.

*Atoh1*-null embryos were received from Dr. Huda Zoghbi at the Baylor College of Medicine. The mice were bred, phenotyped and genotyped by the Zoghbi lab according to the protocol previously described (V. Y. Wang et al., 2005). The animal procedures related to *Atoh1*-null samples are carried out in accordance with The Baylor College of Medicine Institutional Animal Care and Use Committee.

### Tissue preparation and histology

Embryos harvested between E10.5 and E15.5 were immersion-fixed in 4% paraformaldehyde made in 0.1M PB (phosphate buffer) for 1 hour on ice. Embryos harvested at E16.5 and onwards were first trans-cardially perfused with 0.1M PBS (phosphate-buffered saline) and then followed by 4% paraformaldehyde in 0.1M PB, brains were dissected out, and then immersion-fixed in 4% paraformaldehyde in 0.1M PB for 1 hour on ice. Fixed tissues were rinsed thrice with 0.1M PBS followed by cryoprotection in 30% sucrose solution in 0.1M PBS overnight at 4°C before embedding them in optimal cutting temperature (OCT) compound (Tissue-Tek #4583). Tissue was cryosectioned in sagittal orientation at −20°C at 12 µm thickness, mounted on Superfrost slides (Fisher Scientific #12-550-15), air dried at room temperature and stored at - 80°C until use. For all histological experiments, at least 3 different embryos were used with multiple sections per embryo.

### RNA *in situ* hybridization

Digoxigenin-UTP labeled riboprobes (antisense and sense) were generated corresponding to the cDNA of Msx1, Msx2, and Msx3. The cDNA was amplified from a cDNA library made from E12.5 mouse cerebellum using Invitrogen SuperScript IV First-Strand Synthesis System (Invitrogen #18091050). The following primers were used:

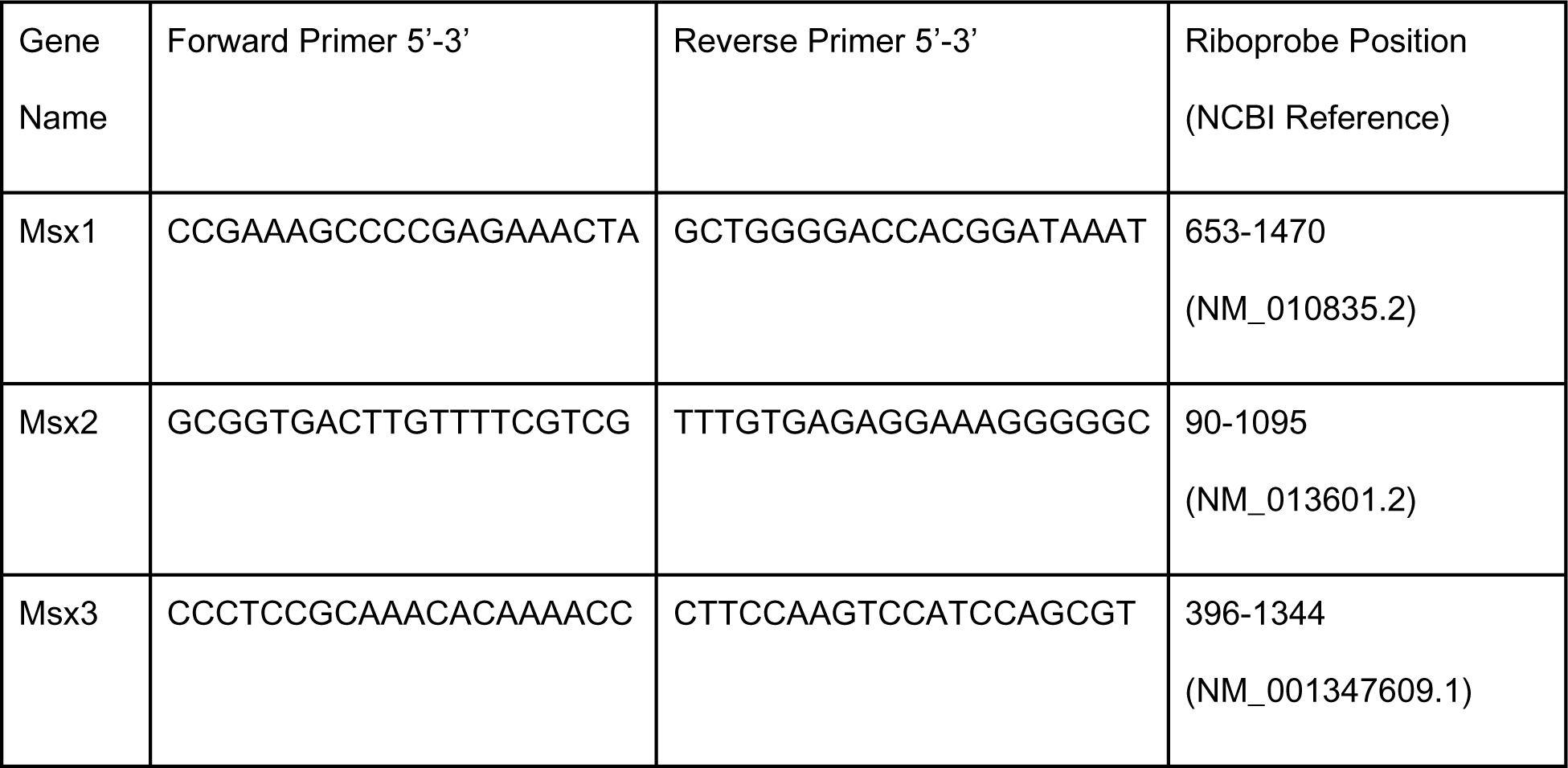

This cDNA was cloned into pGEM-T Easy vector (Promega #A1360) and a combination of gene-specific primers and M13 primers were used to generate DNA templates which were then reverse transcribed using T7 and SP6 polymerases to generate the cRNA probes. The probes were denatured for 10 minutes at 72°C before being added to the hybridization buffer (Ambion, Invitrogen #AM8670). The tissue sections were acetylated with acetic anhydride in 0.1M triethanolamine and dehydrated with increasing concentrations of ethanol before hybridizing them with the probes overnight at 68°C in a humid chamber. Then, they were washed in saline sodium citrate (SSC) solutions: 4xSSC, 2xSSC, 1xSSC and 0.5xSSC at 55°C followed by anti-Digoxigenin antibody (Roche #11093274910, 1:500) incubation for 2 hours at room temperature. After washes, sections were colorized with NBT/BCIP (Roche #11681451001), fixed in 4% PFA, dehydrated and cleared in graded ethanol solutions and Xylene, and coverslipped with Permount diluted in Xylene.

### RNAscope® fluorescence *in situ* hybridization

To look at RNA level expression of 2 genes simultaneously, and at higher resolution, Bio-techne ACD’s RNAscope Multiplex Fluorescent V2 Assay kit (single molecule RNA fluorescent *in situ* hybridization) was used according to manufacturer’s instructions. The RNAscope technology uses tyramide signal amplification which suppresses background and boosts the signal such that individual RNA molecules can be detected as single dot punctae - The “ZZ” probe design only allows amplification to build upon consecutively bound probes on the target, thereby ensuring that each punctate dot represents only real signal (F. Wang et al., 2012; Z. Wang et al., 2013). Briefly, the slides were post-fixed in 4% PFA for 30 minutes, dehydrated in graded ethanol solutions and permeabilized with a protease treatment for 15-30 minutes depending on the tissue age. Slides were then hybridized with the probes overnight at 40°C. After this, the signal amplification tree was built by sequentially incubating slides in Amplifiers 1,2 and 3 at 40°C. The first amplification strand, Amplifier 1, only hybridizes to the “ZZ” s. This was followed by developing the fluorescent channels that involved incubation with HRP attached to the channel-specific sequence, adding the fluorescent dye, and then adding HRP blockers so the other channels can be developed similarly. All these incubations were done at 40°C for durations based on the user manual guide provided by the manufacturer. After the final HRP blocking step, slides were incubated in DAPI to counterstain for 5 minutes before coverslipping with FluorSave mounting medium. RNAscope protocol dictates a short DAPI treatment to ensure that the punctate dots (real signal) are visible and not visually overpowered by the much larger nuclear DAPI staining.

### Immunofluorescence

Tissue sections mounted on slides were warmed on slide warmer at 37°C. Then the sections were rehydrated in 0.1M PBS, followed by incubation in 0.1M PBS-T (0.1% Triton X-100 in 0.1M PBS) for permeabilization. Sections were incubated in a blocking solution (1% BSA and 10% normal serum in 0.1M PBS-T) at room temperature for 30 minutes in a humidified chamber. Subsequently, the blocking solution will be replaced by a diluted primary antibody (see table) in the incubation solution (1% BSA and 5% normal serum in 0.1M PBS-T) and incubated at 4°C overnight in a humidified chamber. The sections were washed 3 times in 0.1PBS-T the next day. The sections were then incubated in secondary antibodies diluted in the incubation solution and counterstained using DAPI. The sections were washed 3 times in 0.1M PB and 1 time in 0.01M PB. Coverslip was applied on the slides with mounting media FluorSave (Millipore #345789).

### BrdU labeling

To look at proliferative cells in the cerebellar ventricular zone, pregnant female mouse with E14.5 embryos was intraperitoneally injected with 5-bromo-deoxyuridine (BrdU, Sigma #B5002; 50 µg/g body weight) 1 hour before collecting the embryos. The pulse duration was 1 hour because the cells are rapidly dividing at this age. Tissue sections were prepared as described above. Tissue sections underwent a 1M HCl incubation at 37°C for 30 minutes post rehydration in 0.1M PBS. The sections were incubated with the Rat anti BrdU primary antibody (1:500, AbCam #AB6326).

### Microscopy

Images were taken using a Zeiss Axiovert 200M microscope with the Axiocam/Axiovision software (Carl Zeiss).

| **Antibody** | **Concentration**<br>**(Immunofluorescence)** | **Source and identifier** |
| --- | --- | --- |
| Rabbit anti-Atoh1 | 1:500 | Proteintech 21215-1-AP |
| Rabbit anti-β-galactosidase | 1:500 | Invitrogen A11132 |
| Rat anti-BrdU | 1:500 | AbCam AB6326 |
| Goat anti-Msx1 | 1:50 | Rndsystems AF5045 |
| Alexa Fluor 594 Chicken anti-Goat IgG (H+L) | 1:1000 | Invitrogen A-21468 |
| Alexa Fluor 488 Chicken anti-Rabbit IgG (H+L) | 1:1000 | Invitrogen A-21441 |
| Alexa Fluor 488 Goat anti-Rat IgG (H+L) | 1:1000 | Jackson ImmunoResearch<br>AB_2338351 |

## Notes

### Competing Interest Statement

The authors have declared no competing interest.

