## Supplemental Figures for "Msx genes delineate a novel molecular map of the developing cerebellar neuroepithelium"

### Supplementary Material:

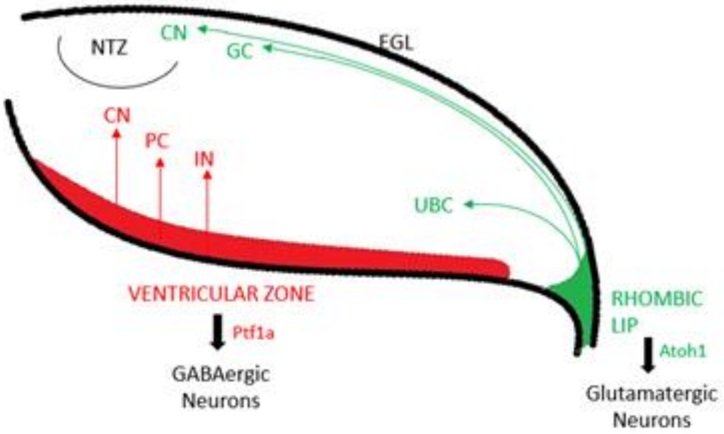


**Supplementary Figure 1. Illustration of the cerebellar progenitor zones in a sagittal view of E12.5 cerebellum.** Cerebellar neurons arise from two primary germinal zones. The glutamatergic lineage comes from the Atoh1 expressing rhombic lip which includes the CN neurons that migrate to the nuclear transitory zone (NTZ), the Granule cells (GC) that form the external granular layer (EGL) and finally the Unipolar Brush cells (UBCs). The GABAergic lineage comes from the Ptf1a expressing ventricular zone that gives rise to the CN neurons, Purkinje cells (PC) and the interneurons (IN). Atoh1 and Ptf1a suppress each other. Illustration as a sagittal section with right side denoting posterior and bottom side ventral.

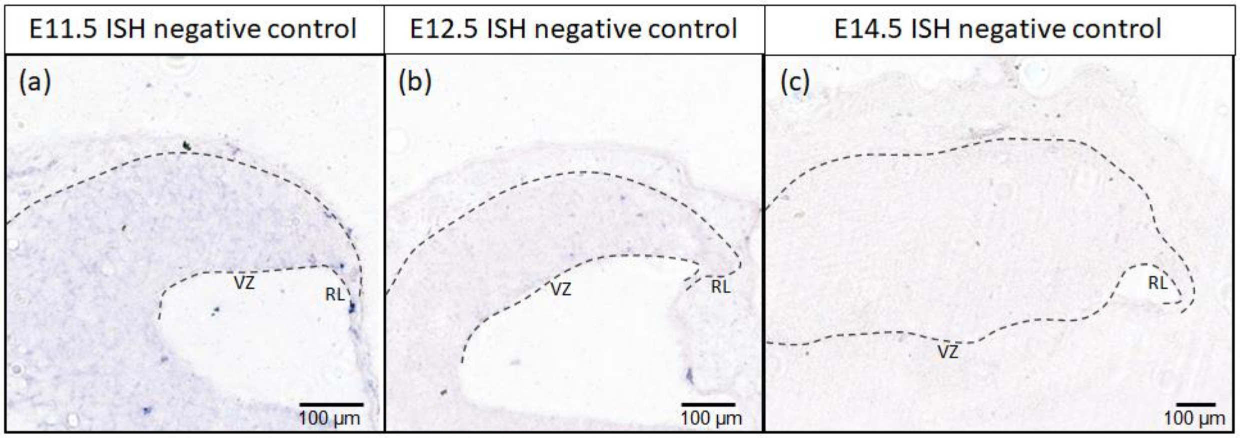


**Supplementary Figure 2. Negative control for RNA in situ hybridization (ISH) for (a) E11.5, (b) E12.5 and (c) E14.5.** Sagittal sections with right side denoting posterior and bottom side ventral. Sense probes of Msx1, Msx2 and Msx3 were combined in equal amounts and used on these sections. RL, rhombic lip; VZ, ventricular zone. Scale bars, 100 µm.

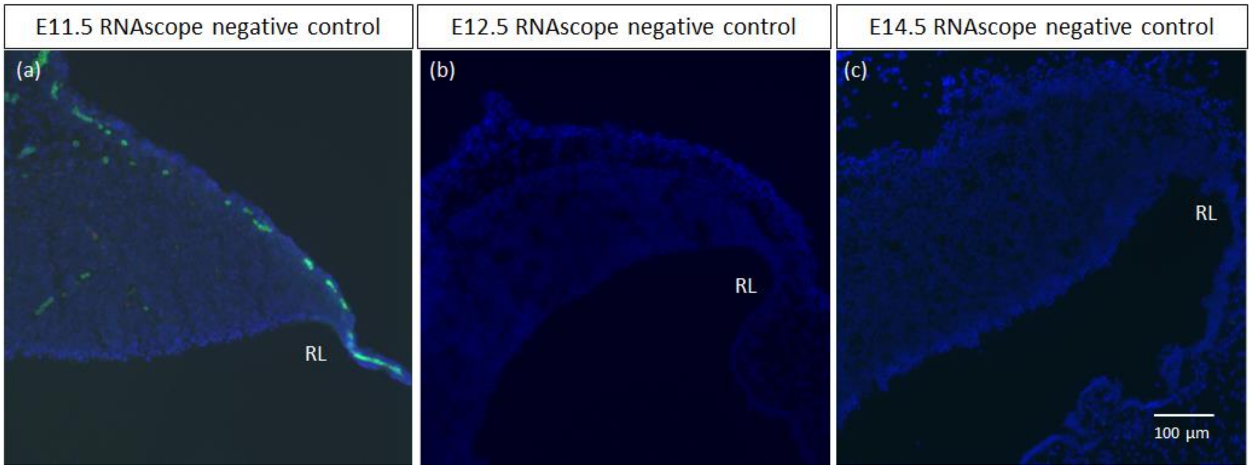


**Supplementary Figure 3. Negative control for RNAscope FISH for (a) E11.5, (b) E12.5 and (c) E14.5.** (a-c) Sagittal sections with right side denoting dorsal and bottom side caudal. Probe for bacterial housekeeping gene (green) was used on these sections, with DAPI (blue) as counterstain. (a) At E11.5 the epithelial roof plate auto-fluoresces to produce the green blob-like artifacts. RL, rhombic lip. Scale bar, 100 µm

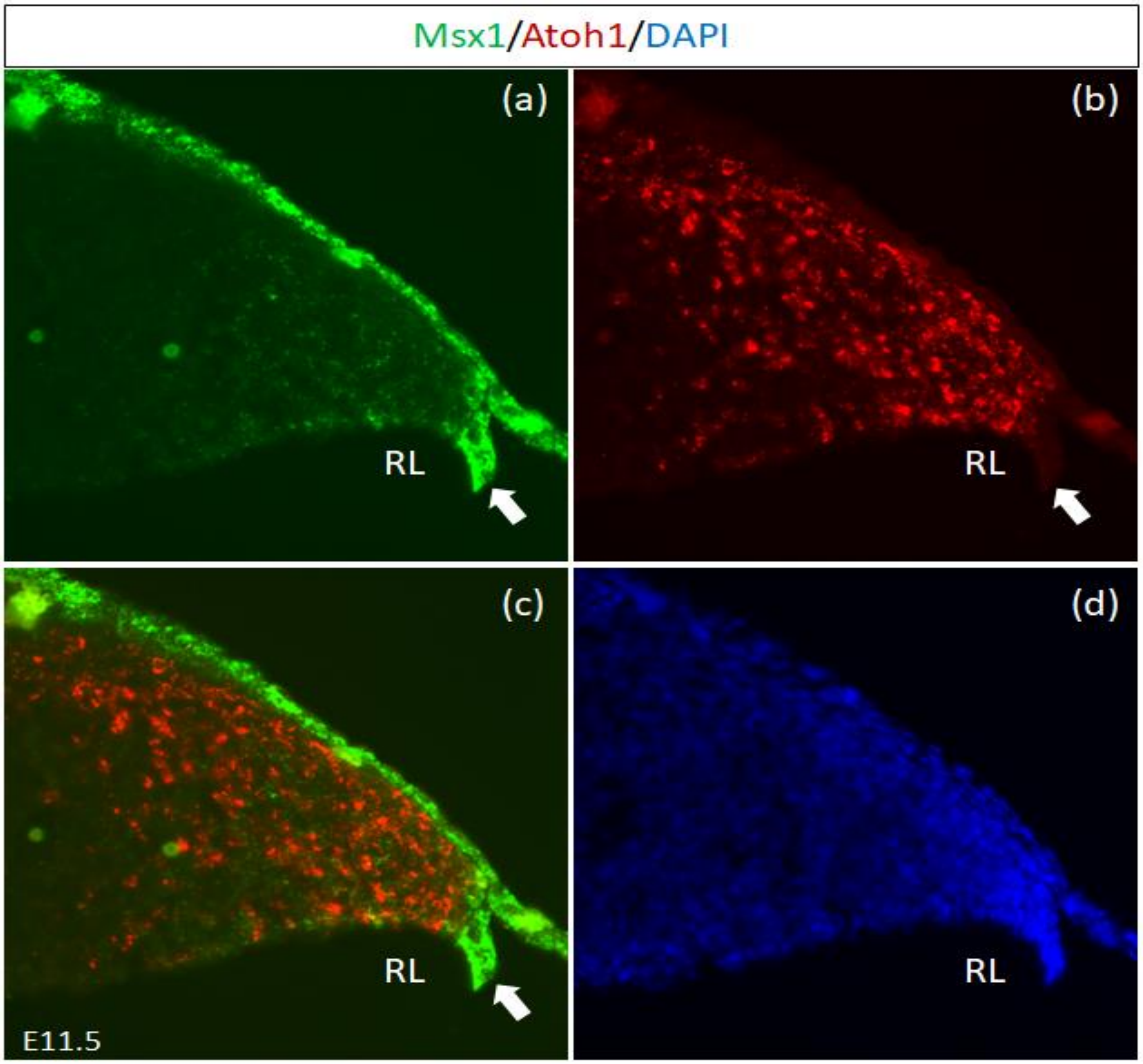


**Supplementary Figure 4. Msx1 and Atoh1 expressions at E11.5**. RNAscope FISH double-label on E11.5 sagittal section. (a) Msx1 (green) is expressed strongest in the caudal-most tip of the RL (white arrows) that is Atoh1 (red) negative (b). (c) Merged Msx1 and Atoh1 staining. (d) DAPI (blue) counterstain for the same tissue section. RL, Rhombic Lip. Scale bar, 100 µm.

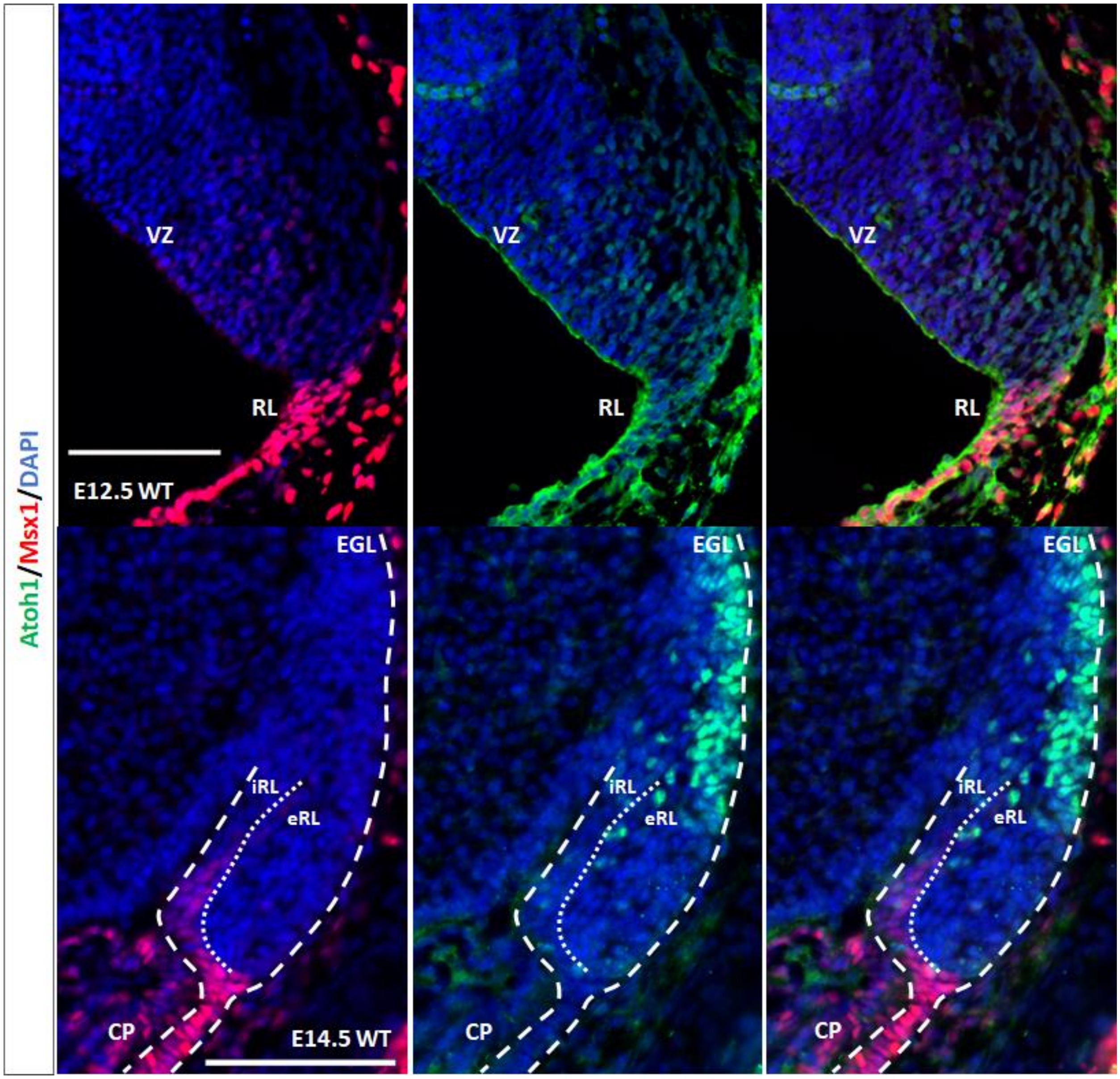


**Supplementary Figure 5. Msx1 and Atoh1 expressions at E12.5 and E14.5**. Immunofluorescence double-label on E12.5 and E14.5 cerebellum. At E12.5, (a) Msx1 (red) is expressed strongest in the caudal-most tip of the RL (white arrows) that is Atoh1 (green) negative (b). (c) Merged Msx1 and Atoh1 staining. At E14.5, Msx1 expression (red in d) is restricted to the interior face of the RL (iRL), which is negative for Atoh1 (green in e). (f) Merged Msx1 and Atoh1 staining illustrated the restricted expression for both molecules at E14.5. CP, choroid plexus; EGL, external germinal layer; RL, Rhombic Lip;, VZ, ventricular zone. Scale bar, 100 µm.

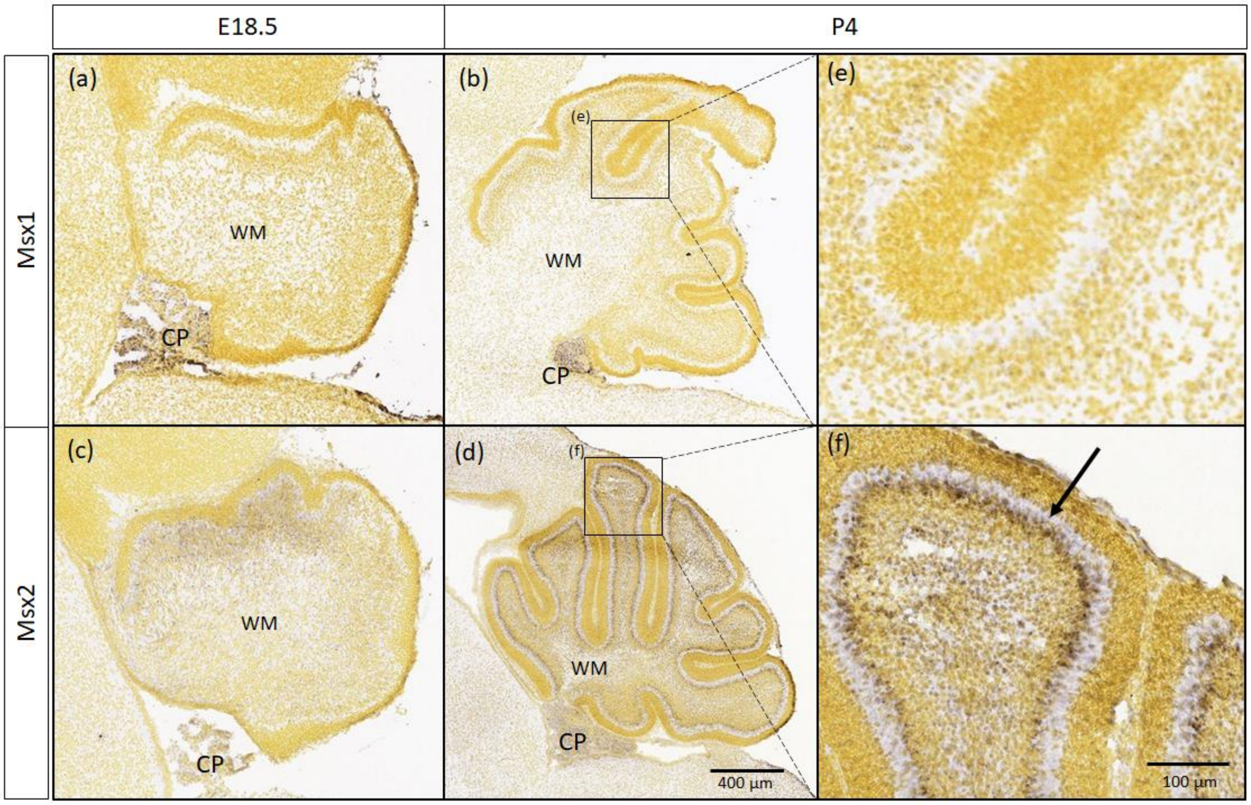


**Supplementary Figure 6. Msx1 and Msx2 expressions in postnatal age.** *In situ* hybridization images taken from the Allen Developing Mouse Brain Atlas (2008). (a-b) Msx1 expression is largely limited to the choroid plexus (CP) and is missing from the cerebellar cortex visible at P4, seen clearly by closeup panel (e). (c-d) Msx2 expression is detected in the developing granule cells as they migrate to form the inner granular layer (IGL) from E18.5 to P4. The Msx2-positive cells in the IGL can be seen clearly in the closeup panel (f) with signal in the IGL (arrow). CP, Choroid Plexus; WM, White Matter. Scale bars indicated 100 µm.

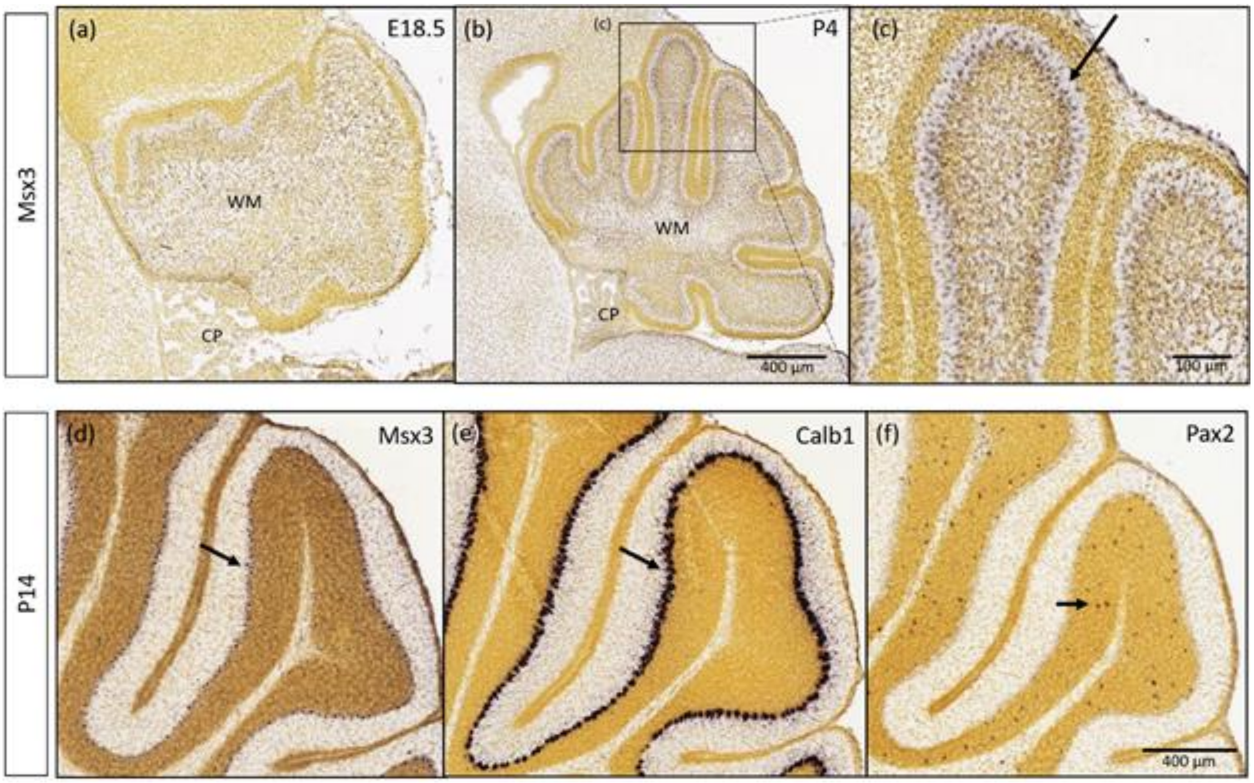


**Supplementary Figure 7. Msx3 expression in postnatal ages localizes to Purkinje cells.** RNA *in situ* hybridization images taken from the Allen Developing Mouse Brain Atlas (2008). (a-c) Msx3 expression is detected in the cerebellum at (a) E18.5 and by (b) P4 Msx3 expression is largely detected in the big cell bodies of the Purkinje cell layer as seen in the closeup panel (c) (arrow). (d-f) By P14, Msx3 expression (d) is clearly visible in the Purkinje cell layer that can be identified by the post-mitotic marker, Calbindin1 at P14 (e) (arrows). (f) shows Pax2-positive post-mitotic interneurons for comparison (arrow). CP, Choroid Plexus; WM, White Matter. Scale bars indicated.
